# High-quality genome of the zoophytophagous stink bug, *Nesidiocoris tenuis*, informs their food habit adaptation

**DOI:** 10.1101/2023.08.29.555439

**Authors:** Tomofumi Shibata, Masami Shimoda, Tetsuya Kobayashi, Hiroshi Arai, Yuta Owashi, Takuya Uehara

## Abstract

The zoophytophagous stink bug, *Nesidiocoris tenuis*, is a promising natural enemy of micro-pests such as whiteflies and thrips. This bug possesses both phytophagous and entomophagous food habits, enabling it to obtain nutrition from both plants and insects. This trait allows us to maintain its population density in agricultural fields by introducing insectary plants, even when the pest prey density is extremely low. However, if the bugs’ population becomes too dense, they can sometimes damage crop plants. This dual character seems to arise from the food preferences and chemosensation of this predator. To understand the genomic landscape of *N. tenuis*, we examined the whole genome sequence of a commercially available Japanese strain. We used long-read sequencing and Hi-C analysis to assemble the genome at the chromosomal level. We then conducted a comparative analysis of the genome with previously reported genomes of phytophagous and hematophagous stink bugs to focus on the genetic factors contributing to this species’ herbivorous and carnivorous tendencies. Our findings suggest that the gustatory gene set plays a pivotal role in adapting to food habits, making it a promising target for selective breeding. Furthermore, we identified the whole genomes of microorganisms symbiotic with this species through genomic analysis. We believe that our results shed light on the food habit adaptations of *N. tenuis* and will accelerate breeding efforts based on new breeding techniques for natural enemy insects, including genomics and genome editing.

## Introduction

Advancements in genome sequencing technology have recently facilitated whole genome sequencing of arthropods, including predatory insects that prey on agricultural pests (Li et al., 2019). However, numerous predatory insects possess minute body sizes and the maintenance of inbred lineages is challenging, resulting in considerable difficulty in obtaining highly contiguous genomes (Leung et al., 2020). By sequencing genomes with longer read length, more comprehensive gene annotation information and polymorphic mutation data can be acquired. Deciphering the whole genome of predatory insects is crucial for the development and selection of superior strains based on genomic information, thereby formulating effective breeding strategies for these natural enemies.

*Nesidiocoris tenuis* (Reuter) (**Fig. 1a**) serves as a predatory insect across extensive regions from Asia to the Mediterranean (Pérez-Hedo and Urbaneja, 2016; Yano, 2022). This species is integral to the integrated pest management of micro-pests. Its predatory activity against pests that pose problems in greenhouse tomato cultivation, such as the silverleaf whitefly (*Bemisia tabaci*) and the tomato leafminer (*Tuta absoluta*), has garnered substantial interest (Calvo et al., 2009; Itou et al., 2013; Sanchez et al., 2009; Urbaneja et al., 2005). Owing to its zoophytophagous nature (**Fig. 1b**), *N. tenuis* can sustain its populations on plant-based food during periods of low pest infestation by incorporating insectary plants and is thus able to mitigate sudden pest outbreaks (Nakano et al., 2021, 2016). However, this insect’s strong preference for insectary plants complicates the control of its movement to crops; moreover, it can cause ring-shaped necrotic damage to tomatoes when the population density becomes excessive (Sanchez et al., 2009; Calvo et al., 2009; Sanchez and Lacasa, 2008). This problem appears to originate from the bug’s feeding habits and chemosensory nature. However, research has focused primarily on behavioral observation (Thomine et al., 2020; Chinchilla-Ramírez et al., 2020), leaving the nature of the insect’s chemosensory receptors and neural substances largely unexplored.

**Figure 1.**
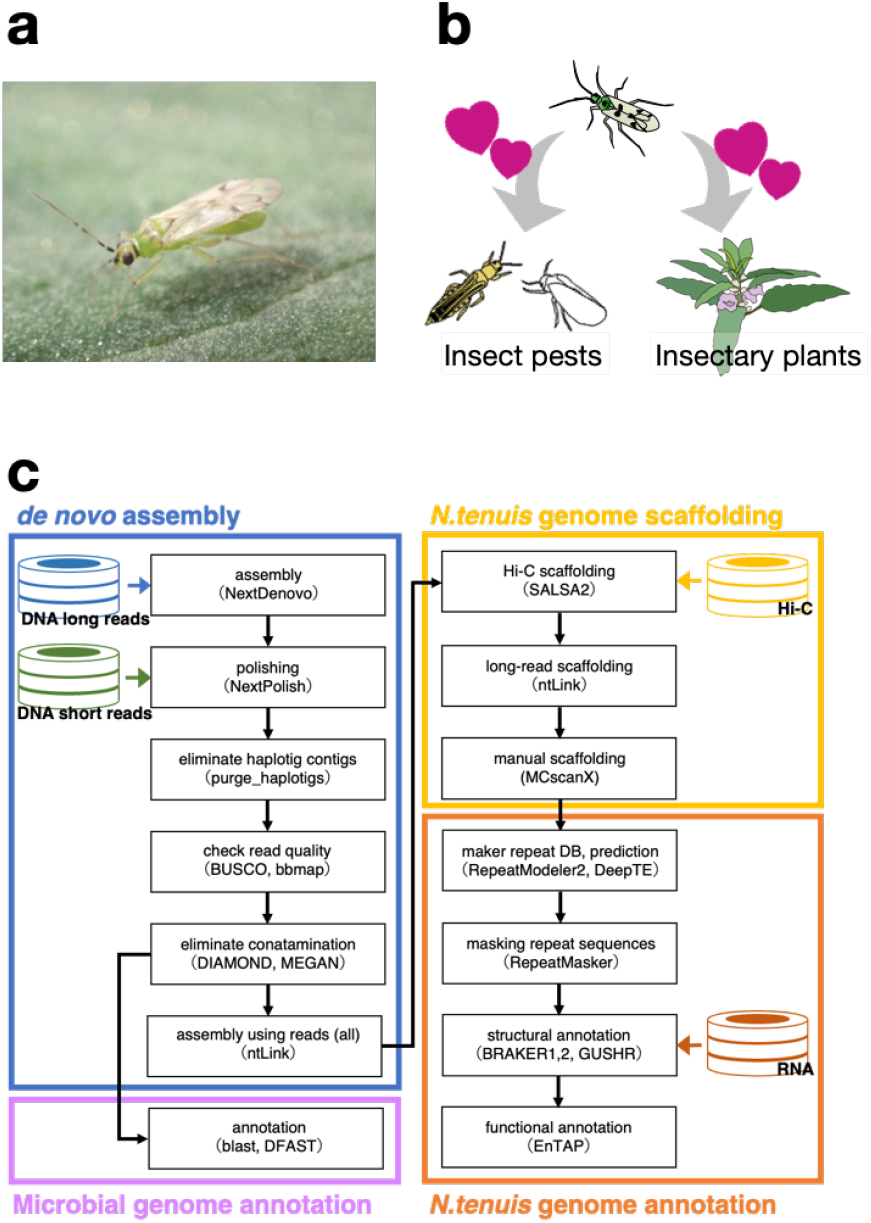
Overviews of the life history of *Nesidiocoris tenuis*and the pipeline used for genome analysis. Adult male of *Nesidiocoris tenuis* on a tomato leaf **(a**), schematic picture of the bug’s zoophytophagy (**b**), and pipeline of genome assembly and analysis (**c**). Cylindrical icons indicate read data and arrows show flow of processed data. The software used and the version in this pipeline are summarized in Supplementary Table 2.

Nevertheless, some pest management approaches that capitalize on these characteristics have been refined through various methods. For instance, the phototaxis of *N. tenuis* can be harnessed to effectively promote movement of the insects from insectary plants to tomato plants (Park and Lee, 2021; Uehara et al., 2019b). Volatiles from infested plants (Rim et al., 2020, 2015, 2018, 2017) and conspecific individuals (Hall et al., 2021) attract *N. tenuis*. Thus, olfactory cues are also favorable candidates for promoting this bug’s movements. With respect to phytophagous feeding habits, research is under way to develop less aggressive strains through selective breeding (Chinchilla-Ramírez et al., 2020) and explore the potential of incorporating RNA interference (RNAi) with pest control strategies (Sarmah et al., 2021). Although a draft genome of this species was published by Ferguson et al. (Ferguson et al., 2021) and the majority of genes have been annotated, further analysis of the genome structure is warranted. In Japan, several cryptic species of *N. tenuis* have been reported (Yasunaga, 2017), necessitating the whole-genome analysis of standard strains employed as biological control agents for the development and implementation of effective integrated pest management strategies.

Here, we investigated the whole genome of a Japanese commercial strain of *N. tenuis* by using long-read sequencing and Hi-C analysis to assemble the genome at the chromosomal level (**Fig. 1c**). A comparative analysis of the genome with the previously reported genomes of phytophagous and hematophagous stink bugs was conducted to elucidate the genetic factors contributing to the herbivorous and carnivorous tendencies of this species. Furthermore, we report on the whole genomes of symbiotic microorganisms of this species identified through genomic analysis. From these results, we discuss feeding habit adaptation through the acquisition of chemosensory genes in *N. tenuis*.

## Results and Discussion

### Genome size and heterozygosity estimates

Short-read sequencing male and female adults produced 26.6 Gb of data. The distribution of *k* -mers using *k* = 21 showed high heterozygosity. We estimated a genome size of 275.9 to 278.3 Mb and a level of heterozygosity of 1.57% to 1.66% (**Supplementary Figure 1**). The genome was about 50 Mb shorter than the previously reported one (Ferguson et al., 2021). In general, heterozygosity tends to be higher in wild insect strains and lower in inbred strains. Our heterozygosity result was not far from those revealed recently for other mirid species’ wild or inbred strain genomes, namely that of *Cyrtorhinus lividipennis* (Bai et al., 2022) was 1.70%; of *Stenotus rubrovittatus* (Kobayashi, 2008), 0.27% to 0.86%, and of *Apolygus lucorum* (Liu et al., 2021), 1%.

### Genome sequencing and assembly

Before performing a genome assembly, we removed low-quality sequences (< Q8) and obtained approximately 29.6 Gb of raw reads (coverage = ×110, **Supplementary Figure 2**). The mean read length and N50 length of filtered reads were 3.4 Kb and 11 Kb, respectively. We performed *de novo* genome assembly with these reads, removed contamination and haplotigs from the assembled genome sequence (**Fig. 1c**), and finally obtained a draft genome with a size of 265.8 Mb. The draft genome had 92 contigs, an N50 of 6.5 Mb, and a GC content of 40%. The draft genome presented here is of high quality compared not only with those of mirid species with high heterozygosity but also with those of other insects with low heterozygosity (**?**Stahlke et al., 2023; Chen et al., 2023).

Subsequently, we obtained a total of 101.5 Gb of raw data from the Hi-C library of the genome and 99.6 Gb of clean reads (without the adapter region sequence). Following chromosome scaffolding and manual curation, we obtained 72 contigs that anchored about 20 chromosomes (**Table 1**). We further selected 17 sequences, starting from the longest to the shortest, and compared their total length to the full genome length. We found that they covered 99.5% of the entire genome. This approach helped us to obtain a complete chromosome-level genome sequence (**Supplementary Figure 3)**. In fact, genomic *in situ* hybridization and comparative genomic hybridization analyses have shown that the karyotype of this species is 2n = 32 (Ferguson et al., 2021). Our result is consistent with this finding. The chromosome-level genome consisted of 16.4 Mb of scaffold N50 and 41.7 Mb of maximum scaffold length. The Benchmarking Universal Single-Copy Orthologue (BUSCO) (Manni et al., 2021) of hemiptera_odb was 94.6% (single copy: 93.6%; duplicated: 1.0%). This chromosome-level assembly provided a substantial increase (>500-fold contiguity) in reference quality compared with the existing reference genome. We mapped raw sequence reads from females to the chromosome-level genome sequence, but there was no coverage bias in each contig; we were therefore unable to determine the sex chromosome.

**Table 1.** Summary statistics of our genomic analysis of *Nesidiocoris tenuis* (NTJ_v2.0) and comparison with the analysis of Ferguson et al. (2021) Adult male of *Nesidiocoris tenuis* on a tomato leaf **(a**), schematic picture of the bug’s zoophytophagy (**b**), and pipeline of genome assembly and analysis (**c**). Cylindrical icons indicate read data and arrows show flow of processed data. The software used and the version in this pipeline are summarized in Supplementary Table 2.

|  | Genome size (Mb) | No. of contigs | No. of scaffolds | Scaffold N50 (Mb) | Scaffold N90 (Mb) | Max scaffold size (Mb) | BUSCO (%) |
| --- | --- | --- | --- | --- | --- | --- | --- |
| NTJ_v2.0 | 265.536 | 72 | 20 | 16.423 | 11.208 | 41.648 | 94.6 |
| Ferguson_v1.5 | 355.120 | 51,852 | 36,513 | 0.287 | 0.033 | 1.392 | 87.9 |

The repeat sequence of this species was 95.9 Mb and about 36.1% of the whole genome sequence (**Fig. 2a**). This ratio is consistent with the ratio estimated from our *k* -mer frequency analysis using GenomeScope 2.0 (see Materials and Methods). The ratios of repeat sequences were 23.7% for Satellite DNA, 11.6% for long tandem repeats, 0.47% for long interspersed elements, and 0.12% for short interspersed elements. Compared with those of the previously reported genome (Ferguson et al., 2021), the long tandem repeat and long interspersed element sequences as a proportion of the whole genome sequences were increased, whereas the ratio of short interspersed elements was decreased. These results suggest that the continuity of the whole-genome sequences was substantially improved.

**Figure 2.**
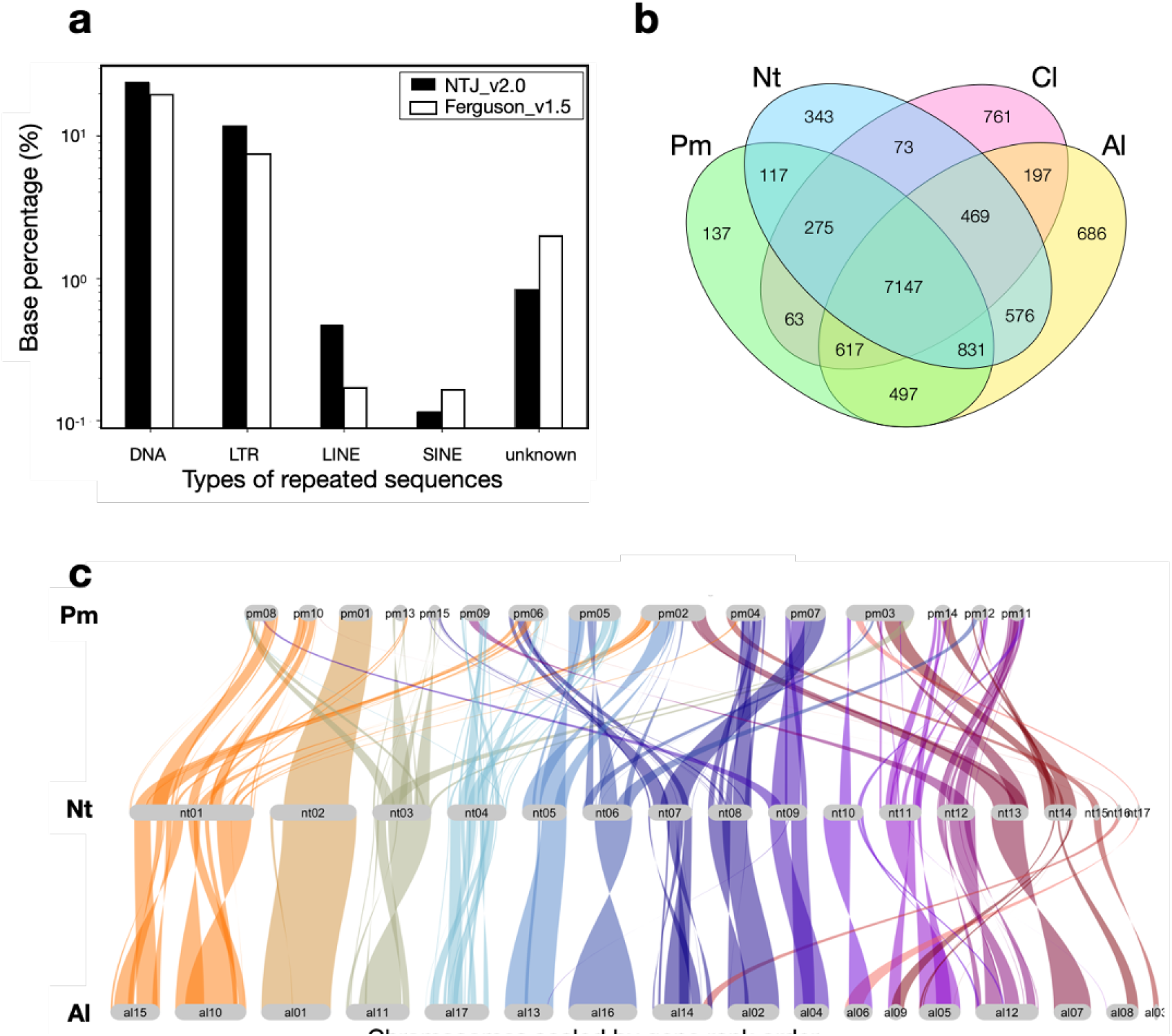
Comparisons of repeated sequences and genome landscapes in hemipteran chromosomes. Comparison of repeated sequences from this study and from that of Ferguson et al. (2021) (**a**), Venn diagram analysis of all annotated genes in *Nesidiocoris tenuis* (Nt) with *Apolygus lucorum* (Al), *Pachypeltis micranthus* (Pm) and *Cimex lectularius* (Cl) (**b**), and synteny analysis of Nt with Al and Pm (**c**).

### Comparative genome analysis of related species

The results of a Venn diagram analysis using the closely related species *A. lucorum* (Liu et al., 2021), *Pachypeltis micranthus* (Yang, 2021), and *Cimex lectularius* (Rosenfeld et al., 2016) by OrthoVenn2 (Xu et al., 2019) showed that *N. tenuis* shared 9023 orthologous genes with *A. lucorum* and 8370 genes with *P. micranthus* (**Fig. 2b**). Additionally, we performed a synteny analysis of the genome NTJ_v2.0 and the chromosomal-level genomes from *A. lucorum* and *P. micranthus* (**Fig. 2c**). The results were consistent with the phylogenetic relatedness among these species. The regions of synteny were about 20% in common reciprocally, suggesting that our chromosomal genome was compatible with the quality of these genomes.

### Complete genomes of symbiotic microorganisms

Generally, antibiotic treatment or removal of the gut is performed to avoid contamination by symbiotic bacterial genomes. However, we were not able to perform this procedure because of the target species’ small body size and the large quantity of genomic DNA (gDNA) required for sequencing. Consequently, we unexpectedly determined the complete genome sequences of symbiotic microorganisms while compiling the draft genome of *N. tenuis*.

On the basis of genome homologies, we confirmed that *N. tenuis* harbored both *Spiroplasma* (Molicutes) and *Rickettsia* (Alphaproteobacteria) endosymbionts, as previously described (Caspi-Fluger et al., 2014; Ferguson et al., 2021; Owashi et al., 2023). The *Spiroplasma* endosymbiont, referred to as the *s*Nten strain, has a main chromosome (1,5 Mb) with six plasmids (1,889 coding sequences in total) and was closely related to the *citri–poulsonii* group (**Fig. 3a**). The *Rickettsia* endosymbiont, referred to as the *r* Nten strain, had a main chromosome (2.3 Mb) with three plasmids (3183 coding sequences in total) and was clustered into the *bellii* group (**Fig. 3b**).

**Figure 3.**
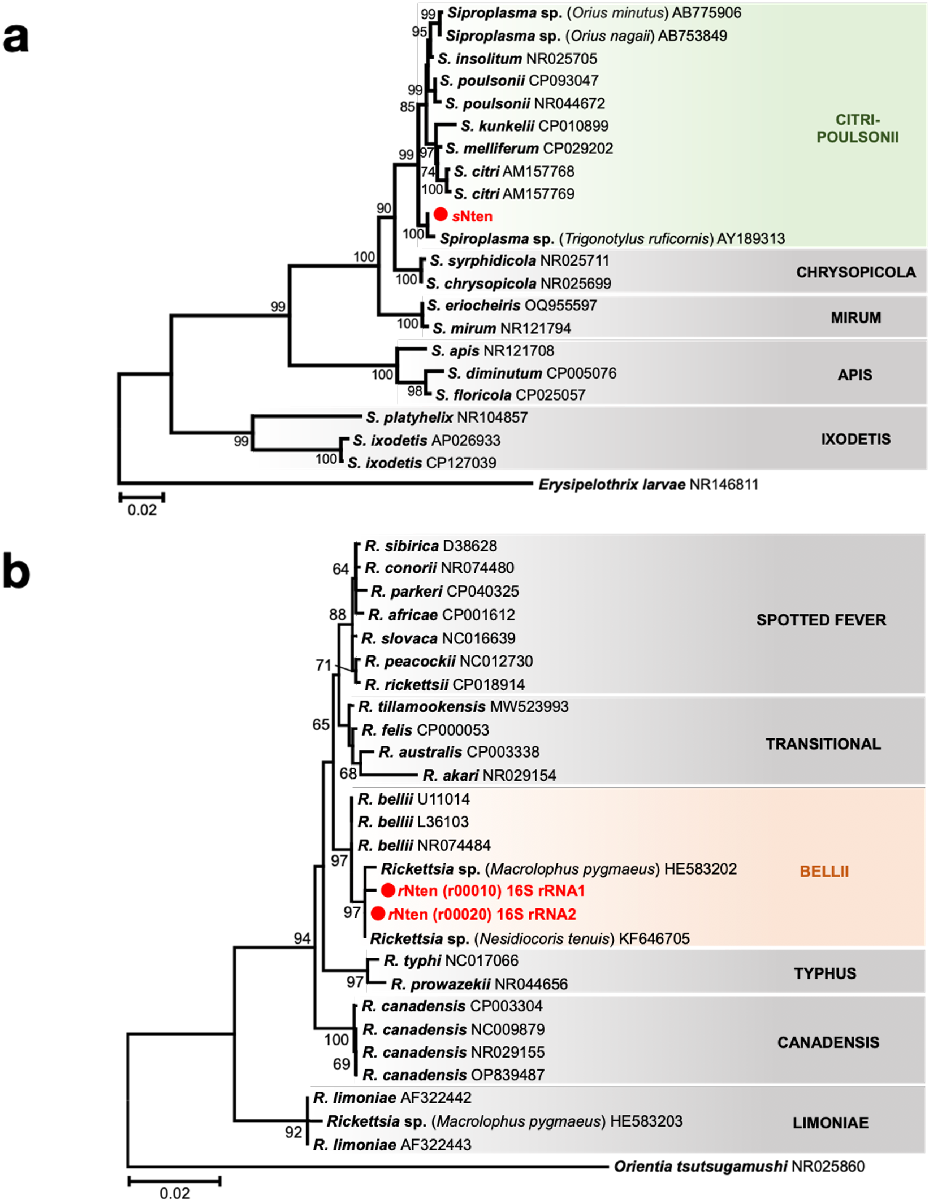
Phylogenetic analysis of symbiotic microorganisms. Phylogenetic trees based on the 16S rRNA genes of *Spiroplasma* and *Rickettsia* were inferred by using the maximum likelihood method on the basis of the Kimura two-parameter model with 1000 bootstrap replicates. Bootstrap values <60% are not shown, and accession numbers are given after each operational taxonomic unit. (**a**) Phylogenetic tree of *Spiroplasma*, based on 1477 positions of the 16S rRNA coding region. The *Spiroplasma* from *Nesidiocoris tenuis* is shown in red. The classification of *Spiroplasma* is represented on the right side and the *citri–poulsonii* clade is shaded green. The outgroup is *Erysipelothrix* larvae and the scale bar indicates 0.02 substitutions per site. (**b**) Phylogenetic tree of *Rickettsia*, based on 1402 positions of the 16S rRNA coding region. The *Rickettsia* from *N. tenuis* is shown in red. The classification of *Rickettsia* is represented on the right side, and the *bellii* clade is shaded with red. The outgroup is *Orientia tsutsugamushi* and the scale bar indicates 0.02 substitutions per site.

*Spiroplasma* relatives of *S. citri* are frequently identified from hemipteran insects (Kwon et al., 1999; Watanabe et al., 2014) and from plants (Saglio et al., 1973; Yokomi et al., 2020). In the case of leafhoppers, *S. citri* is transmitted horizontally between plants and the insect (Mello et al., 2009). In addition, *Spiroplasma* endosymbionts sometimes manipulate their host’s reproduction selfishly in various ways. For example, some *Spiroplasma* strains clustered into the *Spiroplasma ixodetis* and *Spiroplasma poulsonii* groups induce male-killing in their host species, such as ladybirds and fruit flies (Montenegro et al., 2006; Tinsley and Majerus, 2006). In addition, a *citri–poulsonii* group *Spiroplasma* strain induces male-killing in the green lacewing, *Mallada desjardinsi* (Hayashi et al., 2016).

The *Rickettsia* endosymbiont has been detected with high frequency in wild populations of *N. tenuis* in Israel (93% to 100%) and Japan (20.8% to 95.8%) (Caspi-Fluger et al., 2014; Owashi et al., 2023); this is consistent with our detection of *r* Nten in a commercial strain originating from the Japanese wild population. In fact, the nucleotide sequence of *r* Nten was identical to that of the 16S rRNA gene from the Israeli and Japanese populations. In the Israeli *N. tenuis, Rickettsia* were detected in the host gut lumen, suggesting that they may have a nutritional role in the host (Caspi-Fluger et al., 2014). However, the fact that some individuals of wild *N. tenuis* are not infected with *Rickettsia* (Caspi-Fluger et al., 2014; Owashi et al., 2023) means that infection is not obligatory for survival of the host. Some *Rickettsia* endosymbionts induce reproductive phenotypes such as male-killing (Lawson et al., 2001; Schulenburg et al., 2001) and parthenogenesis (Giorgini et al., 2010; Hagimori et al., 2006) in the host.

According to our genome and gene annotation (**Supplementary Table 1**), these genomes have genes encoding enzymes for oxidation or detoxification, besides genes for replicating themselves. We do not have enough data to discuss whether these genes contribute to the host’s feeding habits and survival; further experiments are needed to understand the impact of these symbiotic organisms.

### Chemosensory receptor genes involved in food preference

We constructed a phylogenetic tree with 1286 common single-copy genes from seven hemipteran species, namely *N. tenuis, A. lucorum, Orius laevigatus, C. lectularius, P. micranthus, Rhodnius prolixus*, and *Laodelphax striatellus*. We used *L. striatellus* as an outgroup (**Fig. 4a**). Unlike the other species, *N. tenuis, A. lucorum*, and *P. micranthus* belong to the same family, namely the Miridae. *Pachypeltis micranthus* is phytophagous, whereas *N. tenuis, A. lucorum*, and *O. laevigatus* are zoophytophagous. *C. lectularius, Rhodnius prolixus* and *L. striatellus* are hematophagous. Examination of the phylogenetic tree showed that closely related species were clustered in the same clade, and *A. lucorum* and *P. micranthus* were located closer to each other than to *N. tenuis. Orius laevigatus—*a different genus but also zoophytophagous—was in a different clade from *N. tenuis*, and *R. prolixus* was in the farthest branch. This phylogenetic tree was based on common single-copy genes of these heteropteran insects and therefore reflected the closeness among the species. As mentioned earlier, our Venn diagram comparison of each species with all annotated genes showed that more genes in common were detected between closer species. These results suggest that feeding habits and life histories have diverged but that the ancestral gene sets have not changed greatly, irrespective of the feeding habit.

**Figure 4.**
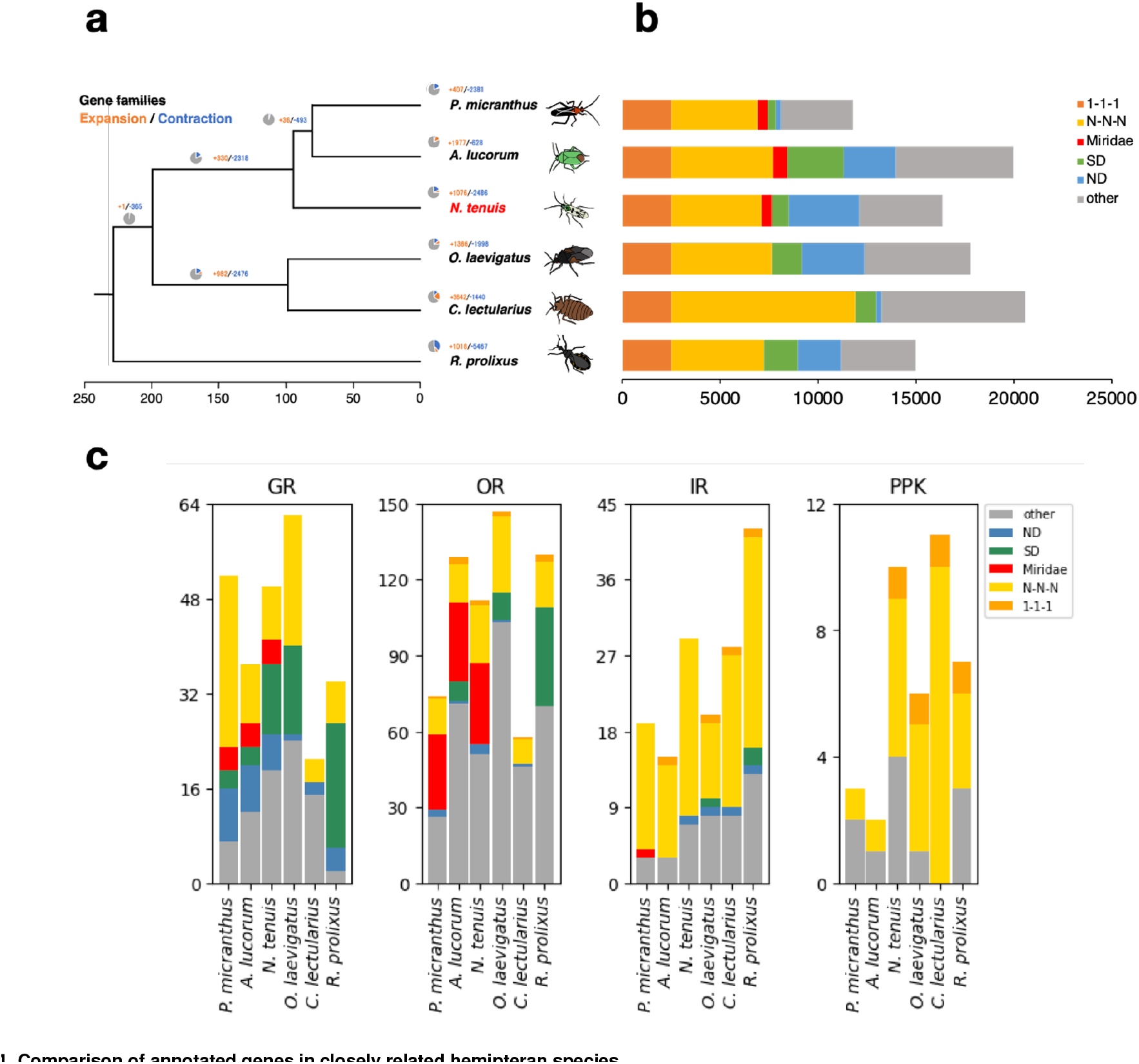
Comparison of annotated genes in closely related hemipteran species. Phylogenetic tree of closely related stink bugs (**a**) and comparison of the number of annotated genes (**b**). Colors representing species in the left panel (**a**) correspond to those in the right panel (**b**). Comparison of chemosensory receptor genes (**c**). Abbreviations are as follows; ND: unclustered genes specific to each species; SD: species-specific genes with multiple gene copies; Miridae: Miridae-genus-specific genes; N-N-N: multicopy universal genes; and 1-1-1: single-copy universal genes.

Divergence and convergence events in all annotated genes were analyzed by using CAFÉ (**Fig. 4a**). We used the CAFÉ analysis to explore gene family expansion and contraction, and we estimated the *N. tenuis* gene birth rate at –0.0416108 with regard to duplications/gene/Mya. We detected 653 gene families that had experienced notable expansion or contraction events across the six species. The results demonstrated that genes had either diverged or converged with speciation, yet we found no universal patterns that mirrored phylogenetic relationships or dietary habits.

We next focused on genes associated with feeding habit, such as chemosensory receptor genes. As chemosensory genes, we focused on olfactory receptor (OR), gustatory receptor (GR), ionotropic receotpr (IR), and *pickpocket* (PPK) of the degenerin/epithelial sodium channel gene family. Chemosensory genes commonly identified in insects, such as olfactory coreceptor (*Orco*), and GRs for sugar, bitter, and CO_2_ sensation were revealed, but we currently lack ligand information or the relevant literature concerning other ORs or GRs. Comparison of gene counts showed characteristic differences between ORs and GRs, while no notable disparities were found in IR and PPK (**Fig. 4b, c**).

For instance, the comparison of OR genes indicated the presence of those specific to Miridae, whereas that of GR genes indicated the existence of those that were more species specific. In fact, our phylogenetic analysis of OR genes frequently revealed mirid-specific and non-mirid-specific OR gene clades throughout the tree. One plausible interpretation for this finding is that ancestral mirid species procured a particular set of OR genes at a certain juncture and, since then, they have utilized similar olfactory signals with a comparable set of ORs (**Fig. 5a**). While OR gene clades reflect phylogenetic relation, these do not align with their food habits. It seems unlikely that ORs play a central role in the adaptation of feeding behaviors. This is consistent with the transition scenario from zoophytophagous to phytophagous habits proposed by Wheeler (Wheeler, 2001).

**Figure 5.**
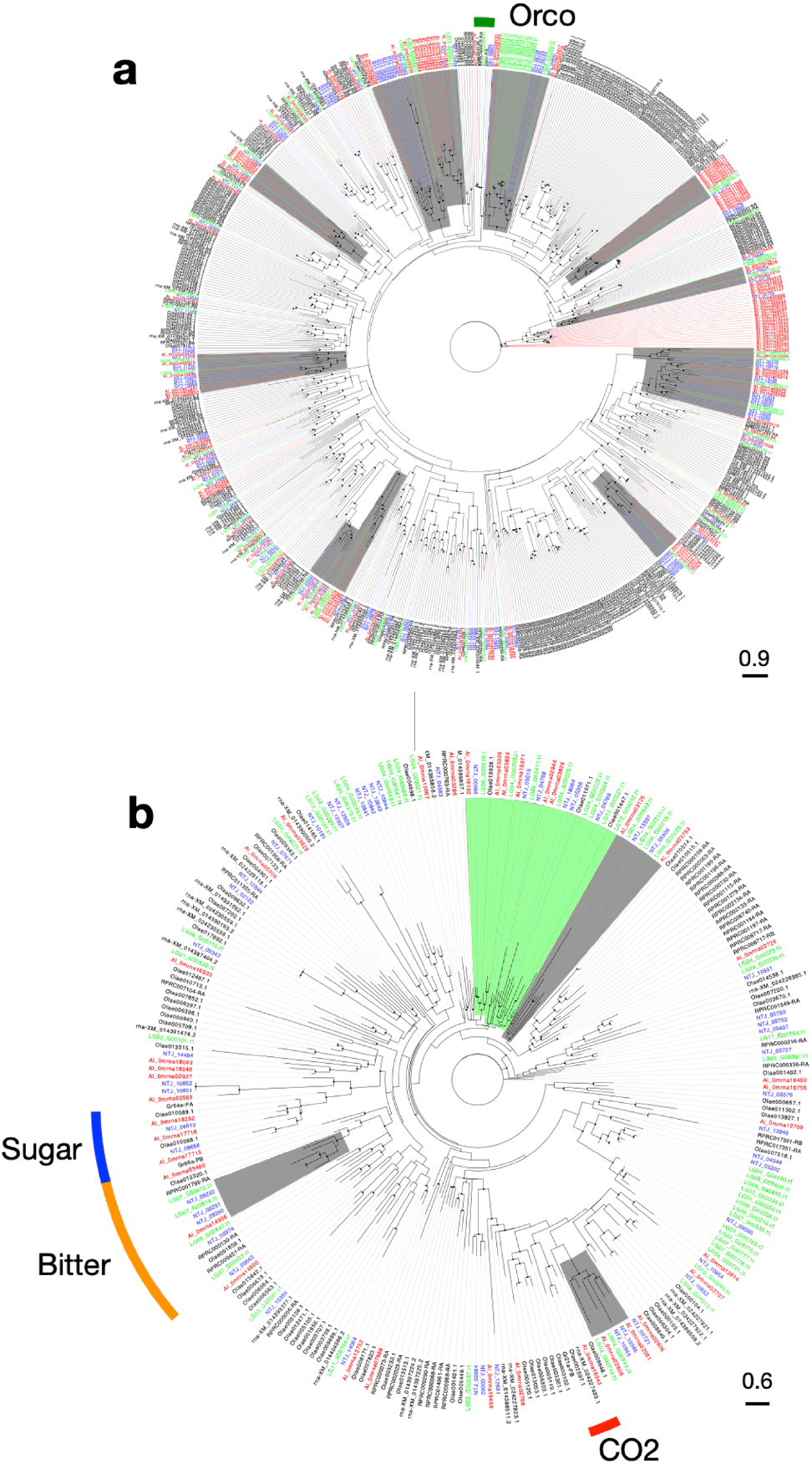
Phylogenetic analysis of chemosensory genes. Phylogenetic trees of olfactory (**a**) and gustatory (**b**) genes. Black circles indicate branches with bootstrap values of 80% or higher. The gray highlights indicate a clade consisting of three species from Miridae (*Apolygus lucorum, Nesidiocoris tenuis, Pachypeltis micranthus*), while the green highlights represent a clade comprising four species with herbivorous feeding habits (A. lucorum, N. tenuis, P. micranthus, *Orius laevigatus*). Each clade is characterized by the presence of at least five genes. Commonly found insect chemosensory receptor genes, such as olfactory coreceptor (*Orco*) and sweet, bitter, and CO2 gustatory receptors are shown.

In contrast, in the case of GR genes, mirid-specific clusters as large as the OR gene clusters were not observed, but were instead identified as small clusters (**Fig. 5b**). Notwithstanding, the herbivore-specific GR genes formed a clade (**Fig. 5b**). These findings suggest that some GRs are common among herbivorous species and may detect similar plant chemicals. However, each species appears to lean more towards acquiring its own specific GRs in alignment with its dietary habits. In terms of the different effective ranges of signals detected by olfaction and gustation, olfactory and gustatory cues generally provide signals for long and close distance, respectively (Bruce et al., 2005). Plants are immobile organisms, so that a combination of these two sensory signals is crucial for finding a host, particularly in herbivorous species (Silva and Clarke, 2020). For example, in lepidopteran insects, the egg-laying preference of female adults determines the larval feeding habitat and survival. It has been suggested that expansion of the availability of the preferred host (i.e., extension of the range of detection of plant secondary metabolites) is linked with the presence of duplicating taste receptors (Engsontia et al., 2014; Suzuki et al., 2018). Our results indicated that the number of GRs was almost identical in zoophytophagous and phytophagous species, yet higher than in hematophagous species. This variation can potentially be elucidated by comparing the diversity of ligand molecules required for detecting animal- and plant-based foods, alongside an examination of the respective nutritional values of each food.

### Future perspectives on breeding natural enemies by using genomics and genome editing

Biological control agents, such as natural enemy insects, represent pest management solutions that incur a small environmental burden and reduce labor intensity. However, as living organisms, these insects may not always exhibit the targeted effectiveness of chemical pesticides. We have developed tools to address this issue thus far (Ogino et al., 2016; Tokushima et al., 2016; Uehara et al., 2019a,b). With the advent of genome editing technology, enhancement of the efficacy of natural enemies through breeding has become increasingly feasible. In this context, we have constructed a highly contiguous genome that can serve not only as a reference sequence for genome editing but also for the analysis of variants in, for example, single nucleotide polymorphisms in field populations. We believe that the genome revealed here will contribute to explorations of the biological functions and beneficial traits of *N. tenuis*.

## Materials and Methods

### Insect

A commercial strain of *N. tenuis* (**Fig. 1a**) was obtained from Agrisect Inc. *Ephestia kuehniella* Zeller eggs (GaRanÒ; Agrisect Inc., Ibaraki, Japan) were used as food, and *Plectranthus amboinicus* (Lamiaceae) was used as an oviposition substrate and moisture source. Bugs were maintained under 25 ± 1 °C, 50% to 70% relative humidity, and a 16:8 light:dark photoperiod. To reduce heterozygosity, we crossed an unmated male and female, placed them together in plastic cases (24 × 17 × 5 cm), and established a sibling line. We used this line for the following sequencing analysis.

### gDNA extraction

High-molecular-weight DNA was extracted by using NucleoBond HMW DNA (MACHEREY-NAGEL GmbH & Co. KG, Düren, Germany) in accordance with the following manufacturers’ protocols. Eight hundred adult males of *N. tenuis* were separated into four groups of 200 individuals each other, and one group was crushed with liquid nitrogen in a mortar and pestle until it became a fine powder. In accordance with the protocol, we obtained 150 μL of final DNA extract. The amount of DNA extracted was adjusted to 9 μg, and this was filtered on the basis of length by using a Short Read Eliminator Kit (Circulomics Inc., MD, USA) to remove any short DNA fragments. A Nanodrop microvolume spectrophotometer (Thermo Fisher Scientific, MA, USA) and a Qubit 4.0 fluorometer (Thermo Fisher Scientific) were used for quality checks on each step of the DNA extraction.

### Library preparation and sequencing

Size-selected gDNA was ligated by using a Ligation Sequencing Kit (SQK-LSK 112, Oxford Nanopore Technologies plc., Oxford, UK) and a NEBNext Companion Module for Oxford Nanopore Technologies Ligation Sequencing (New England Biolabs Inc., MA, USA). It was then purified with Agencourt AMPure XP beads (Beckman Coulter, CA, USA). We followed the protocol of the Ligation Sequencing Kit. One microgram of ligated DNA was sequenced on a Flow Cell (FLO-MIN112, Oxford Nanopore Technologies) by using a MinION sequencer (Oxford Nanopore Technologies) to obtain raw sequence reads.

For the Hi-C analysis, gDNA was extracted from approximately 300 other male adults group that were crushed in liquid nitrogen. A library was prepared by Arima-HiC (Arima Genomics., SD, USA) in accordance with the manufacturer’s protocol. The libraries were sequenced with a Novaseq6000 whole-genome sequencing system.

### Assembly and polishing

The software used is listed in **Supplementary Table 2**. Minion raw data were base-called in high accuracy mode with Guppy v6.0.16. Sequence data were quality checked by using Nanoplot v1.35.5 (De Coster et al., 2018). Genome size and heterozygosity were then estimated by using GenomeScope 2.0 (Ranallo-Benavidez et al., 2020) and paired-end sequences were read on a Novaseq6000 (Illumina Inc.). The estimated genome size was used for genome size specification in the following analyses. Raw reads after quality checking were assembled by using NextDenovo (https://github.com/Nextomics/NextDenovo) with an accurate setting (read_cutoff = 2k). The resulting contigs were polished by using NextPolish v1.3.0 (Hu et al., 2020) with paired-end sequence reads in Novaseq6000 to correct for sequence errors. The resulting genome was verified by using genome statistical values such as N50 (BBTools program stats.sh v44.0, https://jgi.doe.gov/data-and-tools/software-tools/bbtools/) and BUSCO v5.4.5 (Manni et al., 2021). Genome contamination from symbiotic microorganisms was removed by MEGAN and DIAMOND(Bagci et al., 2021). We lastly performed 1-kbp or less raw long-read assembly by using ntLink with the “-gap_fill” option (Coombe et al., 2023).

### Scaffolding

Ligation regions of the Hi-C raw reads were removed by using fastp v0.23.2 (Chen et al., 2018). Scaffolding was performed with the Hi-C reads by using SALSA v2.3 (Ghurye et al., 2019). We next performed raw long-read-based scaffolding by using ntLink (Coombe et al., 2023) and then finally checked manually for remaining gaps. We used GENESPACE (Wang et al., 2012) for the synteny comparison of scaffolded genomes and visualization of the results. The scaffolded genome was also used to create a contact map by using nf-core/hic (Ewels et al., 2020) and visualized by using HiGlass (Kerpedjiev et al., 2018) (Supplementary Fig. 3).

### Structural and functional gene annotation

We identified repeat sequences by using RepeatModeler v2.0.2a (Flynn et al., 2020) to increase the accuracy of the genome structural annotations. Estimated repeat regions were classified by using DeepTE (Yan et al., 2020) and softmasked with RepeatMasker v4.1.2.p1 (Tarailo-Graovac and Chen, 2009). The soft-masked genome was subjected to the gene prediction software BRAKER1 (Hoff et al., 2016) and BRAKER2 (Bruna et al., 2021). The predicted region was integrated by using TSEBRA v1.0.3 (Gabriel et al., 2021), and then untranslated region sequences were added by using GUSHR v1.0.0 (Keilwagen et al., 2016). For functional annotation, we used EnTAP v0.5.0 (Hart et al., 2020) with databases such as SwissProt or TrEMBL for invertebrates.

The genome data (contigs) of the symbiotic microbes were annotated with BLAST searches and DFAST (Tanizawa et al., 2017). Circularity of the contigs was confirmed by using BLASTN searches, and each contig was closed manually as described by Arai et al. (Arai et al., 2022).

### Phylogenetic and comparative analysis of genes

For phylogenetic analysis, we chose six hemipteran species with different feeding habits, namely *A. lucorum* (Liu et al., 2021), *O. laevigatus* (Bailey et al., 2022), *C. lectularius* (Rosenfeld et al., 2016), *P. micranthus* (Gäde and Marco, 2022), *R. prolixus* (Mesquita et al., 2015), and *L. striatellus* (Zhu et al., 2017). We obtained the protein sequences of the six species by using gff-read v0.12.7 (Pertea and Pertea, 2020) from genome (FASTA) and annotation (GTF; gene transfer format) data, and we removed pseudogenes by using seqkit v2.3.1 (Shen et al., 2016). We then chose a single copy core gene set in the BUSCO database (hemiptera_-odb10) from the protein sequences and constructed an interspecific phylogenetic tree by using iqtree v2.2.0.3 (Minh et al., 2020). The phylogenetic tree was converted to a tree considering divergence age by using r8s v1.81 (Sanderson, 2003). Genes orthologous among the six species were classified into orthogroups by OrthoFinder v2.5.4 (Emms and Kelly, 2019). We estimated gene gain and loss rates by using CAFE v4.2.1 (Han et al., 2013) from divergence, considering the phylogenetic tree and orthogroups.

We chose chemosensory receptors (GR, OR, IR, PKK), which have typical protein motif structures, from the Pfam database. Receptor genes were then aligned by using mafft v7.508 (Nakamura et al., 2018), the alignment was trimmed with trimal v1.4.1 (Capella-Gutierrez et al., 2009), the phylogenetic tree was created with iqtree v2.2.0.3, and visualization was performed with Figtree v1.4.4 (https://github.com/rambaut/figtree).

The phylogenetic trees of the symbiotic microorganisms were drawn from the 16S rRNA gene inferred by using the maximum likelihood method based on the Kimura two-parameter model with 1000 bootstrap replicates. Bootstrap values <60% are not shown and accession numbers are given after each operational taxonomic unit.

## Data availability

All raw reads obtained for genome assembly have been deposited in the DNA Data Bank of Japan under Bio-ProjectID PRJDB16217 and Sequence Read Archive (SRA) under DRR495435.

## Acknowledgments

We thank Dr. Toshiyuki Tezuka of Agrisect inc. for providing insects colony. We also acknowledge the efforts of the members of the Insect Design Technology Group for maintenance of the colony. We are grateful to the members of the Laboratory of Applied Entomology at the University of Tokyo for their invaluable comments on this study. This project was supported by JSPS KAKENHI (#20H02992); JSPS Fostering Joint International Research (#21KK0273); and in part by the Cabinet Office, Government of Japan, Cross-ministerial Moonshot Agriculture, Forestry and Fisheries Research and Development Program (#JPJ009237). This research was in part the fulfillment of an MSc degree (by TS) at the University of Tokyo.

## Contributions

T.S. and T.U. conceived the project and designed and interpreted all of the experiments. M.S. and T.K. helped to design and perform insect husbandry and sample preparations. T.S. and T.U. performed experiments and collected data. T.S. assembled, curated, and analyzed the genome data. Y.O. and H.A. analyzed the whole genome of symbiotic microorganisms. T.S. and T.U. wrote the manuscript.

## Supplementary Information

**Figure S1.**
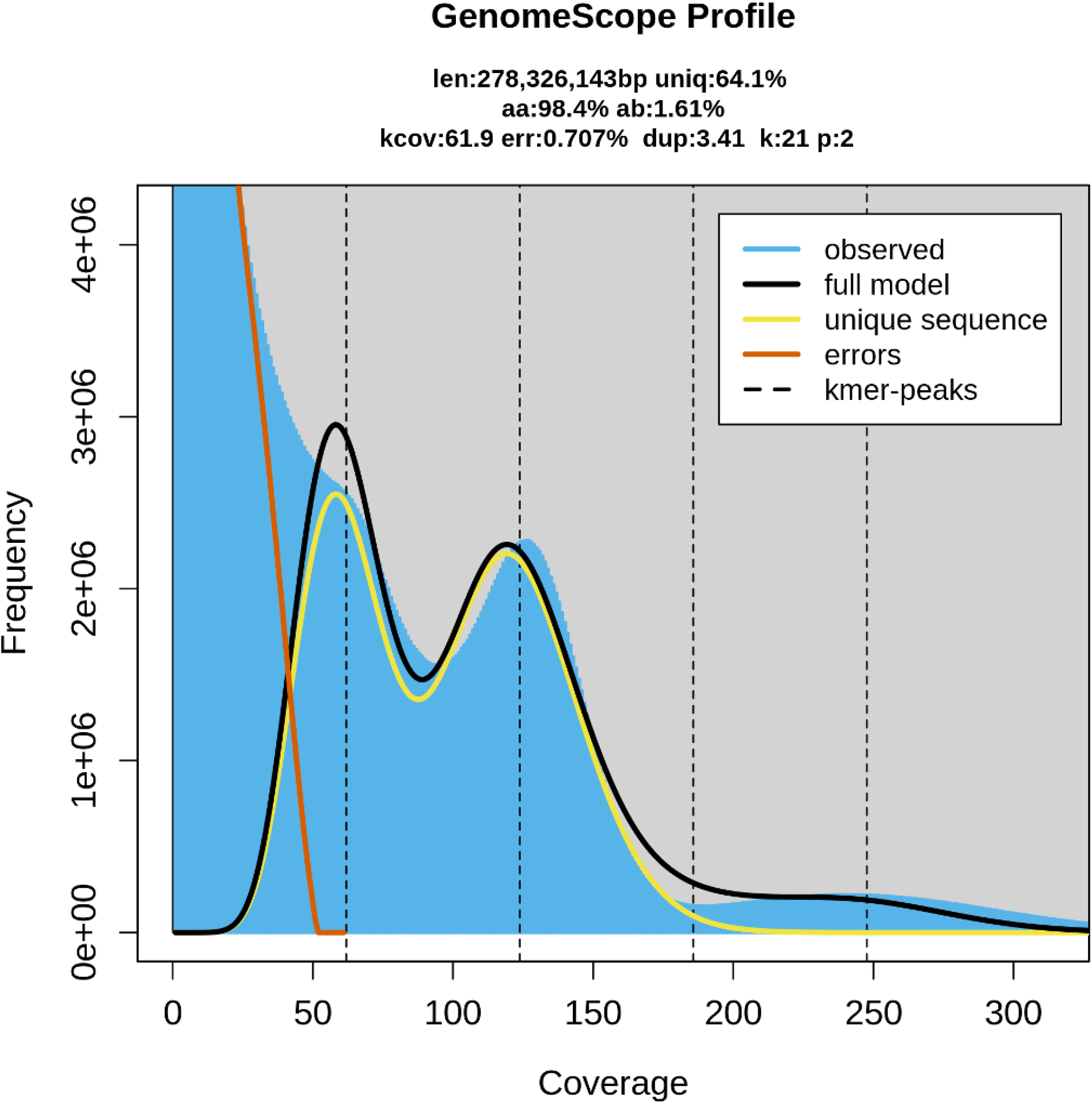
GenomeScope 2.0 profile predicts genome size as 278Mb and heterogeneity as 1.61%.

**Figure S2.**
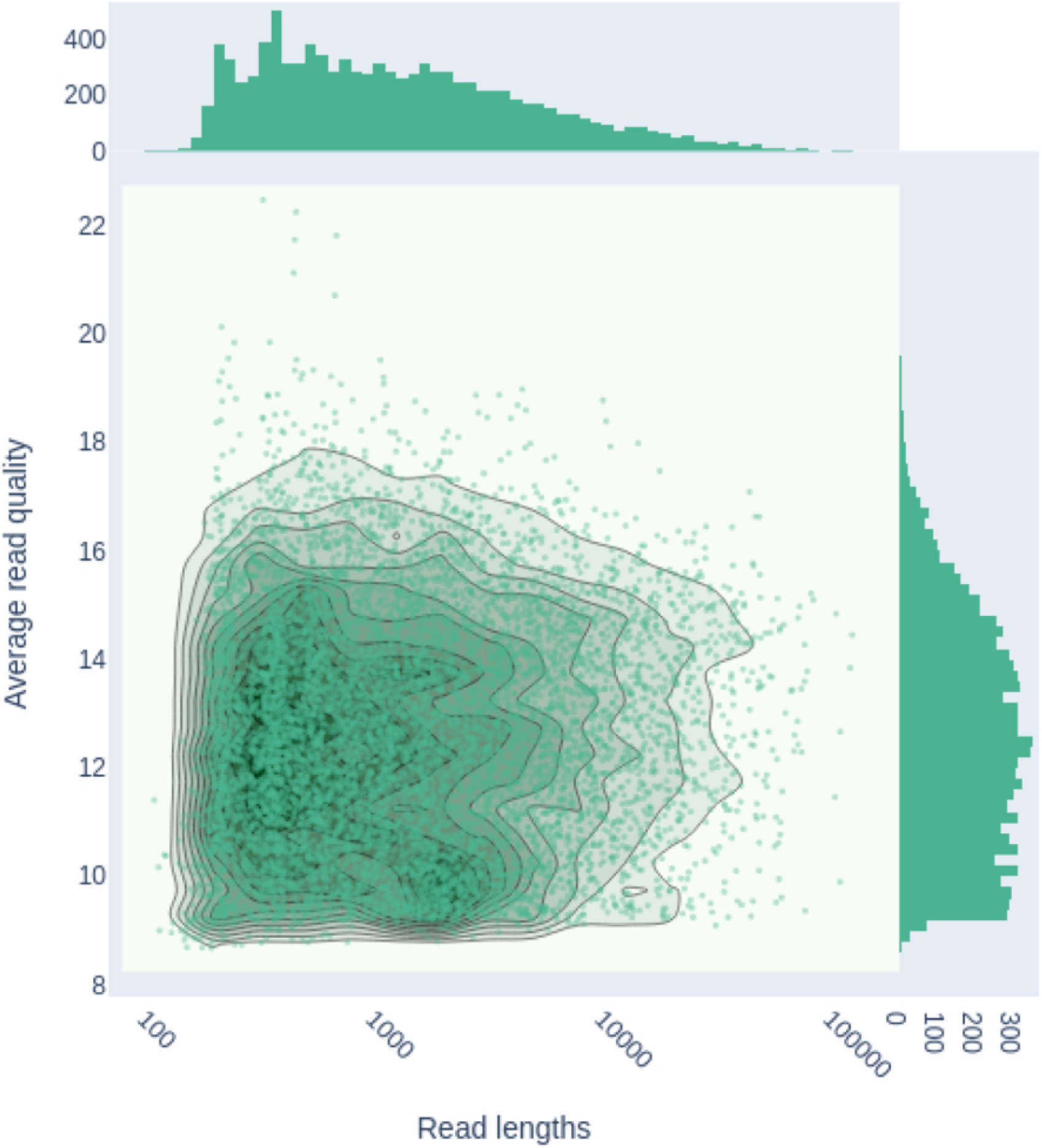
Quality map of nanopore sequencing reads.

**Figure S3.**
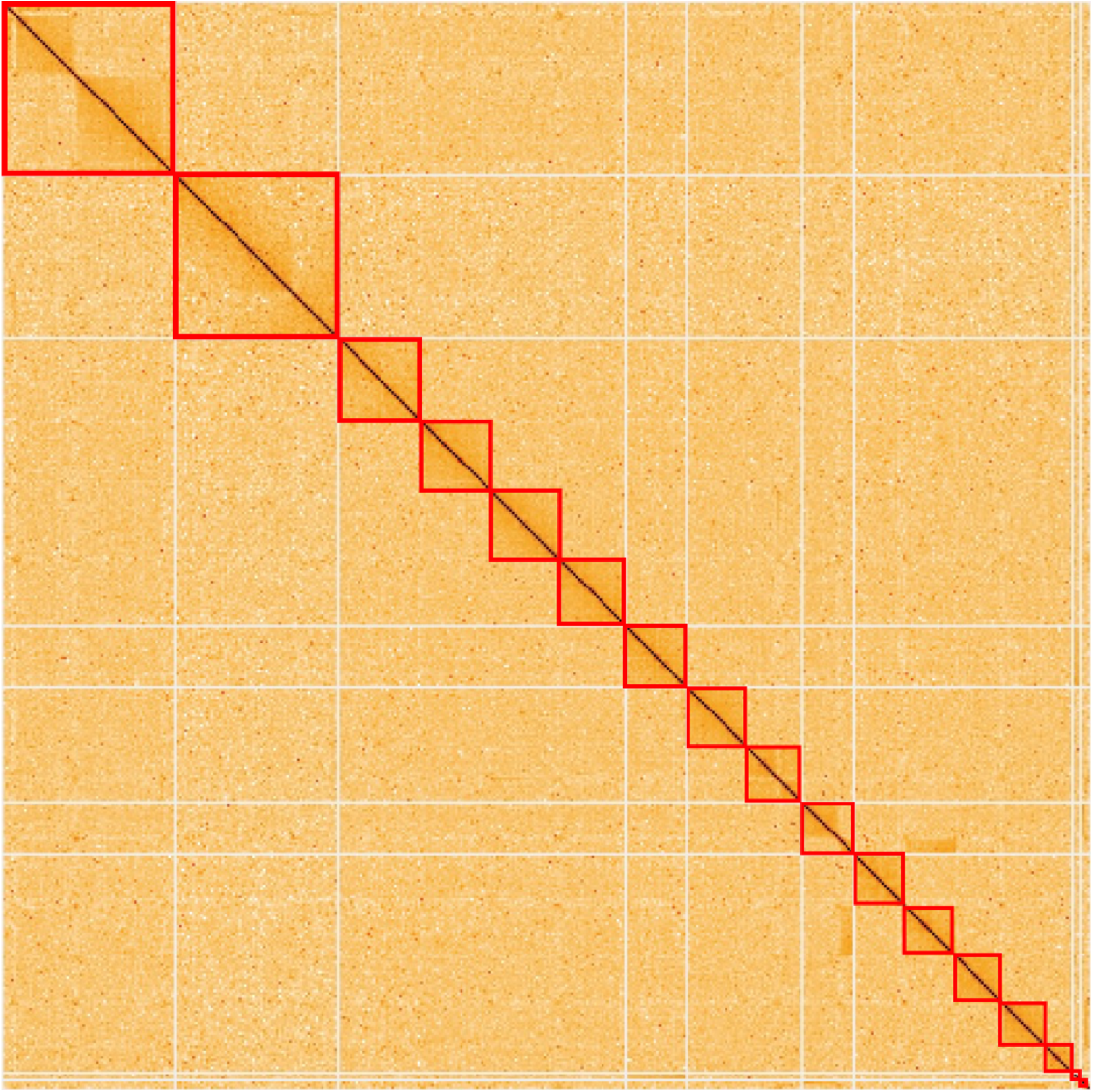
Contact map from Hi-C data suggests 17 chromosomes.

## Notes

### Competing Interest Statement

The authors have declared no competing interest.

## Reference

Arai, H., Inoue, M. N., and Kageyama, D. 2022. Male-killing mechanisms vary between spiroplasma species. Frontiers in Microbiology, 13:1075199.

Bagci, C., Patz, S., and Huson, D. H. 2021. Diamond+megan: Fast and easy taxonomic and functional analysis of short and long microbiome sequences. Curr Protoc, 1(3):e59.

Bai, Y., Shi, Z., Zhou, W., Wang, G., Shi, X., He, K., Li, F., and Zhu, Z. R. 2022. Chromosome-level genome assembly of the mirid predator cyrtorhinus lividipennis reuter (hemiptera: Miridae), an important natural enemy in the rice ecosystem. Mol Ecol Resour, 22(3):1086–1099.

Bailey, E., Field, L., Rawlings, C., King, R., Mohareb, F., Pak, K. H., Hughes, D., Williamson, M., Ganko, E., Buer, B., and Nauen, R. 2022. A scaffold-level genome assembly of a minute pirate bug, orius laevigatus (hemiptera: Anthocoridae), and a comparative analysis of insecticide resistance-related gene families with hemipteran crop pests. BMC Genomics, 23(1):45.

Bruce, T. J., Wadhams, L. J., and Woodcock, C. M. 2005. Insect host location: a volatile situation. Trends in plant science, 10(6):269–274.

Bruna, T., Hoff, K. J., Lomsadze, A., Stanke, M., and Borodovsky, M. 2021. Braker2: automatic eukaryotic genome annotation with genemark-ep+ and augustus supported by a protein database. NAR Genom Bioinform, 3(1):qaa108.

Calvo, J., Bolckmans, K., Stansly, P. A., and Urbaneja, A. 2009. Predation by nesidiocoris tenuis on bemisia tabaci and injury to tomato. BioControl, 54(2):237–246.

Capella-Gutierrez, S., Silla-Martinez, J. M., and Gabaldon, T. 2009. trimal: a tool for automated alignment trimming in large-scale phylogenetic analyses. Bioinformatics, 25 (15):1972–1973.

Caspi-Fluger, A., Inbar, M., Steinberg, S., Friedmann, Y., Freund, M., Mozes-Daube, N., and Zchori-Fein, E. 2014. Characterization of the symbiont rickettsia in the mirid bug nesidiocoris tenuis (reuter) (heteroptera: Miridae). Bulletin of Entomological Research, 104(6):681–688.

Chen, L., Yu, X.-Y., Xue, X.-F., Zhang, F., Guo, L.-X., Zhang, H.-M., Hoffmann, A. A., Hong, X.-Y., and Sun, J.-T. 2023. The genome sequence of a spider mite, tetranychus truncatus, provides insights into interspecific host range variation and the genetic basis of adaptation to a low-quality host plant. Insect Science.

Chen, S., Zhou, Y., Chen, Y., and Gu, J. 2018. fastp: an ultra-fast all-in-one fastq preprocessor. Bioinformatics, 34(17):i884–i890.

Chinchilla-Ramírez, M., Pérez-Hedo, M., Pannebakker, B. A., and Urbaneja, A. 2020. Genetic variation in the feeding behavior of isofemale lines of nesidiocoris tenuis. Insects, 11(8):513.

Coombe, L., Warren, R. L., Wong, J., and Nikolic, V. 2023. ntlink: a toolkit for de novo genome assembly scaffolding and mapping using long reads. arXiv.

De Coster, W., D’Hert, S., Schultz, D. T., Cruts, M., and Van Broeckhoven, C. 2018. Nanopack: visualizing and processing long-read sequencing data. Bioinformatics, 34 (15):2666–2669.

Emms, D. M. and Kelly, S. 2019. Orthofinder: phylogenetic orthology inference for comparative genomics. Genome Biol, 20(1):238.

Engsontia, P., Sangket, U., Chotigeat, W., and Satasook, C. 2014. Molecular evolution of the odorant and gustatory receptor genes in lepidopteran insects: Implications for their adaptation and speciation. Journal of Molecular Evolution, 79(1):21–39.

Ewels, P. A., Peltzer, A., Fillinger, S., Patel, H., Alneberg, J., Wilm, A., Garcia, M. U., Di Tommaso, P., and Nahnsen, S. 2020. The nf-core framework for community-curated bioinformatics pipelines. Nature biotechnology, 38(3):276–278.

Ferguson, K. B., Visser, S., Dalíková, M., Provazníková, I., Urbaneja, A., Pérez-Hedo, M., Marec, F., Werren, J. H., Zwaan, B. J., and Pannebakker, B. A. 2021. Jekyll or hyde? the genome (and more) of nesidiocoris tenuis, a zoophytophagous predatory bug that is both a biological control agent and a pest. Insect Molecular Biology, 30(2):188–209.

Flynn, J. M., Hubley, R., Goubert, C., Rosen, J., Clark, A. G., Feschotte, C., and Smit, A. F. 2020. Repeatmodeler2 for automated genomic discovery of transposable element families. Proc Natl Acad Sci U S A, 117(17):9451–9457.

Gabriel, L., Hoff, K. J., Bruna, T., Borodovsky, M., and Stanke, M. 2021. Tsebra: transcript selector for braker. BMC Bioinformatics, 22(1):566.

Ghurye, J., Rhie, A., Walenz, B. P., Schmitt, A., Selvaraj, S., Pop, M., Phillippy, A. M., and Koren, S. 2019. Integrating hi-c links with assembly graphs for chromosome-scale assembly. PLoS Comput Biol, 15(8):e1007273.

Giorgini, M., Bernardo, U., Monti, M. M., Nappo, A. G., and Gebiola, M. 2010. Rickettsia symbionts cause parthenogenetic reproduction in the parasitoid wasp pnigalio soemius (hymenoptera: Eulophidae). Applied and Environmental Microbiology, 76(8):2589–2599.

Gäde, G. and Marco, H. G. 2022. The adipokinetic peptides of hemiptera: Structure, function, and evolutionary trends. Frontiers in Insect Science, 2.

Hagimori, T., Abe, Y., Date, S., and Miura, K. 2006. The first finding of a rickettsia bacterium associated with parthenogenesis induction among insects. Current Microbiology, 52(2): 97–101.

Hall, D. R., Harte, S. J., Bray, D. P., Farman, D. I., James, R., Silva, C. X., and Fountain, M. T. 2021. Hero turned villain: Identification of components of the sex pheromone of the tomato bug, nesidiocoris tenuis. Journal of Chemical Ecology, 47:394–405.

Han, M. V., Thomas, G. W., Lugo-Martinez, J., and Hahn, M. W. 2013. Estimating gene gain and loss rates in the presence of error in genome assembly and annotation using cafe 3. Mol Biol Evol, 30(8):1987–1997.

Hart, A. J., Ginzburg, S., Xu, M. S., Fisher, C. R., Rahmatpour, N., Mitton, J. B., Paul, R., and Wegrzyn, J. L. 2020. Entap: Bringing faster and smarter functional annotation to non-model eukaryotic transcriptomes. Mol Ecol Resour, 20(2):591–604.

Hayashi, M., Watanabe, M., Yukuhiro, F., Nomura, M., and Kageyama, D. 2016. A nightmare for males? a maternally transmitted male-killing bacterium and strong female bias in a green lacewing population. PLoS One, 11(6):e0155794.

Hoff, K. J., Lange, S., Lomsadze, A., Borodovsky, M., and Stanke, M. 2016. Braker1: Unsupervised rna-seq-based genome annotation with genemark-et and augustus. Bioinformatics, 32(5):767–769.

Hu, J., Fan, J., Sun, Z., and Liu, S. 2020. Nextpolish: a fast and efficient genome polishing tool for long-read assembly. Bioinformatics, 36(7):2253–2255.

Itou, M., Watanabe, M., Watanabe, E., and Miura, K. 2013. Gut content analysis to study predatory efficacy of nesidiocoris tenuis (reuter) (hemiptera: Miridae) by molecular methods. Entomological Science, 16(2):145–150.

Keilwagen, J., Wenk, M., Erickson, J. L., Schattat, M. H., Grau, J., and Hartung, F. 2016. Using intron position conservation for homology-based gene prediction. Nucleic Acids Research, 44(9):e89–e89.

Kerpedjiev, P., Abdennur, N., Lekschas, F., McCallum, C., Dinkla, K., Strobelt, H., Luber, J. M., Ouellette, S. B., Azhir, A., Kumar, N., et al. (2018). Higlass: web-based visual exploration and analysis of genome interaction maps. Genome biology, 19(1):1–12.

Kobayashi, T. 2008. Development of polymorphic microsatellite markers for the sorghum plant bug, stenotus rubrovittatus (heteroptera: Miridae). Mol Ecol Resour, 8(3):690–691.

Kwon, M.-O., Wayadande, A. C., and Fletcher, J. 1999. Spiroplasma citri movement into the intestines and salivary glands of its leafhopper vector, circulifer tenellus. Phytopathology®, 89(12):1144–1151.

Lawson, E. T., Mousseau, T. A., Klaper, R., Hunter, M. D., and Werren, J. H. 2001. Rickettsia associated with male-killing in a buprestid beetle. Heredity, 86(4):497–505.

Lee, J., Nishiyama, T., Shigenobu, S., Yamaguchi, K., Suzuki, Y., Shimada, T., Katsuma, S., and Kiuchi, T. 2021. The genome sequence of samia ricini, a new model species of lepidopteran insect. Molecular ecology resources, 21(1):327–339.

Leung, K., Ras, E., Ferguson, K. B., Ariëns, S., Babendreier, D., Bijma, P., Bourtzis, K., Brodeur, J., Bruins, M. A., Centurión, A., Chattington, S. R., Chinchilla-Ramírez, M., Dicke, M., Fatouros, N. E., González-Cabrera, J., Groot, T. V. M., Haye, T., Knapp, M., Koskinioti, P., …, and Pannebakker, B. A. 2020. Next-generation biological control: the need for integrating genetics and genomics. Biological Reviews, 95(6):1838–1854.

Li, F., Zhao, X., Li, M., He, K., Huang, C., Zhou, Y., Li, Z., and Walters, J. R. 2019. Insect genomes: progress and challenges. Insect Molecular Biology, 28(6):739–758.

Liu, Y., Liu, H., Wang, H., Huang, T., Liu, B., Yang, B., Yin, L., Li, B., Zhang, Y., Zhang, S., Jiang, F., Zhang, X., Ren, Y., Wang, B., Wang, S., Lu, Y., Wu, K., Fan, W., and Wang, G. 2021. Apolygus lucorum genome provides insights into omnivorousness and mesophyll feeding. Mol Ecol Resour, 21(1):287–300.

Manni, M., Berkeley, M. R., Seppey, M., Simao, F. A., and Zdobnov, E. M. 2021. Busco update: Novel and streamlined workflows along with broader and deeper phylogenetic coverage for scoring of eukaryotic, prokaryotic, and viral genomes. Mol Biol Evol, 38(10): 4647–4654.

Mello, A. F. S., Wayadande, A. C., Yokomi, R. K., and Fletcher, J. 2009. Transmission of different isolates of spiroplasma citri to carrot and citrus by circulifer tenellus (hemiptera: Cicadellidae). Journal of Economic Entomology, 102(4):1417–1422.

Mesquita, R. D., Vionette-Amaral, R. J., Lowenberger, C., Rivera-Pomar, R., Monteiro, F. A., Minx, P., Spieth, J., Carvalho, A. B., Panzera, F., Lawson, D., Torres, A. Q., Ribeiro, J. M., Sorgine, M. H., Waterhouse, R. M., Montague, M. J., Abad-Franch, F., Alves-Bezerra, M., and Oliveira, P. L. 2015. Genome of Rhodnius prolixus, an insect vector of Chagas disease, reveals unique adaptations to hematophagy and parasite infection. Proc Natl Acad Sci U S A, 112(48):14936–14941.

Minh, B. Q., Schmidt, H. A., Chernomor, O., Schrempf, D., Woodhams, M. D., von Haeseler, A., and Lanfear, R. 2020. Iq-tree 2: New models and efficient methods for phylogenetic inference in the genomic era. Mol Biol Evol, 37(5):1530–1534.

Montenegro, H., Petherwick, A. S., Hurst, G. D. D., and Klaczko, L. B. 2006. Fitness effects of wolbachia and spiroplasma in drosophila melanogaster. Genetica, 127(1):207–215.

Nakamura, T., Yamada, K. D., Tomii, K., and Katoh, K. 2018. Parallelization of mafft for large-scale multiple sequence alignments. Bioinformatics, 34(14):2490–2492.

Nakano, R., Tsuchida, Y., Ishikawa, R., Tatara, A., Amano, Y., Muramatsu, Y., et al. (2016). Control of bemisia tabaci (gennadius) on tomato in greenhouses by a combination of nesidiocoris tenuis (reuter) and banker plants. Annual Report of the Kansai Plant Protection Society, pages 65–72.

Nakano, R., Morita, T., Okamoto, Y., Fujiwara, A., Yamanaka, T., and Adachi-Hagimori, T. 2021. Cleome hassleriana plants fully support the development and reproduction of nesidiocoris tenuis. BioControl, 66:407–418.

Ogino, T., Uehara, T., Muraji, M., Yamaguchi, T., Ichihashi, T., Suzuki, T., Kainoh, Y., and Shimoda, M. 2016. Violet led light enhances the recruitment of a thrip predator in open fields. Scientific reports, 6(1):32302.

Kainoh, and Shimoda]Owashi2023 Owashi, Y., Minami, T., Kikuchi, T., Yoshida, A., Nakano, R., Kageyama, D., and Adachi-Hagimori, T. 2023. Microbiome of zoophytophagous biological control agent Nesidiocoris tenuis. Microbial Ecology

Park, Y. G. and Lee, J. H. 2021. Uv-led lights enhance the establishment and biological control efficacy of nesidiocoris tenuis (reuter) (hemiptera: Miridae). PLoS One, 16(1): e0245165.

Pertea, G. and Pertea, M. 2020. Gff utilities: Gffread and gffcompare. F1000Res, 9.

Pérez-Hedo, M. and Urbaneja, A. 2016. The zoophytophagous predator nesidiocoris tenuis: a successful but controversial biocontrol agent in tomato crops. Advances in insect control and resistance management, pages 121–138.

Ranallo-Benavidez, T. R., Jaron, K. S., and Schatz, M. C. 2020. Genomescope 2.0 and smudgeplot for reference-free profiling of polyploid genomes. Nat Commun, 11(1):1432.

Rim, H., Uefune, M., Ozawa, R., and Takabayashi, J. 2015. Olfactory response of the omnivorous mirid bug nesidiocoris tenuis to eggplants infested by prey: specificity in prey developmental stages and prey species. Biological Control, 91:47–54.

Rim, H., Uefune, M., Ozawa, R., Yoneya, K., and Takabayashi, J. 2017. Experience of plant infestation by the omnivorous arthropod nesidiocoris tenuis affects its subsequent responses to prey-infested plant volatiles. BioControl, 62:233–242.

Rim, H., Uefune, M., Ozawa, R., and Takabayashi, J. 2018. An omnivorous arthropod, nesidiocoris tenuis, induces gender-specific plant volatiles to which conspecific males and females respond differently. Arthropod-Plant Interactions, 12:495–503.

Rim, H., Hattori, S., and Arimura, G.-i. 2020. Mint companion plants enhance the attraction of the generalist predator nesidiocoris tenuis according to its experiences of conspecific mint volatiles. Scientific reports, 10(1):2078.

Rosenfeld, J. A., Reeves, D., Brugler, M. R., Narechania, A., Simon, S., Durrett, R., Foox, J., Shianna, K., Schatz, M. C., Gandara, J., Afshinnekoo, E., Lam, E. T., Hastie, A. R., Chan, S., Cao, H., Saghbini, M., Kentsis, A., Planet, P. J., Kholodovych, V., and Mason, C. E. 2016. Genome assembly and geospatial phylogenomics of the bed bug cimex lectularius. Nat Commun, 7:10164.

Saglio, P., Lhospital, M., Laflèche, D., Dupont, G., Bové, J. M., Tully, J. G., and Freundt, E. A. 1973. Spiroplasma citri gen. and sp. n.: A mycoplasma-like organism associated with “stubborn” disease of citrus. International Journal of Systematic and Evolutionary Microbiology, 23(3):191–204.

Sanchez, J. A. and Lacasa, A. 2008. Impact of the zoophytophagous plant bug nesidiocoris tenuis (heteroptera: Miridae) on tomato yield. Journal of Economic Entomology, 101(6): 1864–1870.

Sanchez, J. A., Lacasa, A., Arnó, J., Castañé, C., and Alomar, O. 2009. Life history parameters for nesidiocoris tenuis (reuter) (het., miridae) under different temperature regimes. Journal of Applied Entomology, 133(2):125–132.

Sanderson, M. J. 2003. r8s: inferring absolute rates of molecular evolution and divergence times in the absence of a molecular clock. Bioinformatics, 19(2):301–302.

Sarmah, N., Kaldis, A., Taning, C. N. T., Perdikis, D., Smagghe, G., and Voloudakis, A. 2021. dsrna-mediated pest management of tuta absoluta is compatible with its biological control agent nesidiocoris tenuis. Insects, 12(4):274.

Schulenburg, J. H. G. v. d., Habig, M., Sloggett, J. J., Webberley, K. M., Bertrand, D., Hurst, G. D. D., and Majerus, M. E. N. 2001. Incidence of male-killing rickettsia spp. (α-proteobacteria) in the ten-spot ladybird beetle adalia decempunctata l. (coleoptera: Coccinellidae). Applied and Environmental Microbiology, 67(1):270–277.

Shen, W., Le, S., Li, Y., and Hu, F. 2016. Seqkit: A cross-platform and ultrafast toolkit for fasta/q file manipulation. PLoS One, 11(10):e0163962.

Silva, R. and Clarke, A. R. 2020. The “sequential cues hypothesis”: a conceptual model to explain host location and ranking by polyphagous herbivores. Insect Science, 27(6): 1136–1147.

Stahlke, A. R., Chang, J., Tembrock, L. R., Sim, S. B., Chudalayandi, S., Geib, S. M., Scheffler, B. E., Perera, O. P., Gilligan, T. M., Childers, A. K., et al. (2023). A chromosome-scale genome assembly of a helicoverpa zea strain resistant to bacillus thuringiensis cry1ac insecticidal protein. Genome Biology and Evolution, 15(3):evac131.

Suzuki, H. C., Ozaki, K., Makino, T., Uchiyama, H., Yajima, S., and Kawata, M. 2018. Evolution of gustatory receptor gene family provides insights into adaptation to diverse host plants in nymphalid butterflies. Genome Biology and Evolution, 10(6):1351–1362.

Tanizawa, Y., Fujisawa, T., and Nakamura, Y. 2017. Dfast: a flexible prokaryotic genome annotation pipeline for faster genome publication. Bioinformatics, 34(6):1037–1039.

Tarailo-Graovac, M. and Chen, N. 2009. Using repeatmasker to identify repetitive elements in genomic sequences. Current Protocols in Bioinformatics, 25(1):4.10.11–14.10.14.

Thomine, E., Jeavons, E., Rusch, A., Bearez, P., and Desneux, N. 2020. Effect of crop diversity on predation activity and population dynamics of the mirid predator nesidiocoris tenuis. Journal of Pest Science, 93:1255–1265.

Tinsley, M. C. and Majerus, M. E. N. 2006. A new male-killing parasitism: Spiroplasma bacteria infect the ladybird beetle anisosticta novemdecimpunctata (coleoptera: Coccinellidae). Parasitology, 132(6):757–765.

Tokushima, Y., Uehara, T., Yamaguchi, T., Arikawa, K., Kainoh, Y., and Shimoda, M. 2016. Broadband photoreceptors are involved in violet light preference in the parasitoid fly exorista japonica. PLoS One, 11(8):e0160441.

Uehara, T., Maeda, T., Shimoda, M., Fujiwara-Tsujii, N., and Yasui, H. 2019. Identification and characterization of the pheromones in the minute pirate bug orius sauteri (heteroptera: Anthocoridae). Journal of Chemical Ecology, 45(10):811–817.

Uehara, T., Ogino, T., Nakano, A., Tezuka, T., Yamaguchi, T., Kainoh, Y., and Shimoda, M. 2019. Violet light is the most effective wavelength for recruiting the predatory bug nesidiocoris tenuis. BioControl, 64(2):139–147.

Urbaneja, A., Tapia, G., and Stansly, P. 2005. Influence of host plant and prey availability on developmental time and survival of nesidiocoris tenius (het.: Miridae). Biocontrol Science and Technology, 15(5):513–518.

Wang, Y., Tang, H., Debarry, J. D., Tan, X., Li, J., Wang, X., Lee, T. H., Jin, H., Marler, B., Guo, H., Kissinger, J. C., and Paterson, A. H. 2012. Mcscanx: a toolkit for detection and evolutionary analysis of gene synteny and collinearity. Nucleic Acids Res, 40(7):e49.

Watanabe, M., Yukuhiro, F., Maeda, T., Miura, K., and Kageyama, D. 2014. Novel strain of spiroplasma found in flower bugs of the genus orius (hemiptera: Anthocoridae): Transovarial transmission, coexistence with wolbachia and varied population density. Microbial Ecology, 67(1):219–228.

Wheeler, A. G. Biology of the plant bugs (Hemiptera: Miridae): pests, predators, opportunists. Cornell University Press, 2001.

Xu, L., Dong, Z., Fang, L., Luo, Y., Wei, Z., Guo, H., Zhang, G., Gu, Y. Q., Coleman-Derr, D., Xia, Q., et al. (2019). Orthovenn2: a web server for whole-genome comparison and annotation of orthologous clusters across multiple species. Nucleic acids research, 47 (W1):W52–W58.

Yan, H., Bombarely, A., and Li, S. 2020. Deepte: a computational method for de novo classification of transposons with convolutional neural network. Bioinformatics, 36(15): 4269–4275.

Yang, B. 2021. Genome sequencing of pachypeltis micranthus mu et liu (hemiptera: Miridae), a potential biological control agent for mikania micrantha. doi: 10.5061/dryad.0k6djhb18.

Yano, E. 2022. Biological control using zoophytophagous bugs in japan. Journal of Pest Science, 95(4):1473–1484.

Yasunaga, T. 2017. Three new species of nesidiocoris kirkaldy from japan, with a taxonomic review of the tribe dicyphini for eastern asia (heteroptera: Miridae: Bryocorinae). Tijdschrift voor Entomologie, 160(1):25–40.

Yokomi, R., Chen, J., Rattner, R., Selvaraj, V., Maheshwari, Y., Osman, F., Pagliaccia, D., and Vidalakis, G. 2020. Genome sequence resource for spiroplasma citri, strain cc-2, associated with citrus stubborn disease in california. Phytopathology, 110(2):254–256.

Zhu, J., Jiang, F., Wang, X., Yang, P., Bao, Y., Zhao, W., Wang, W., Lu, H., Wang, Q., Cui, N., Li, J., Chen, X., Luo, L., Yu, J., Kang, L., and Cui, F. 2017. Genome sequence of the small brown planthopper, laodelphax striatellus. Gigascience, 6(12):1–12.

